# Changes in Extracellular Matrix Gene and Protein Expressions in Human Trabecular Meshwork Cells in Response to Mechanical Fluid Flow Stimulation

**DOI:** 10.1101/796094

**Authors:** Koichi Yoshida, Motofumi Kawai, Tsugiaki Utsunomiya, Akihiro Ishibazawa, Young-Seok Song, Mariana Sayuri B. Udo, Yoshikazu Tasaki, Akitoshi Yoshida

**Affiliations:** Department of Ophthalmology, Asahikawa Medical University, Asahikawa, Hokkaido, Japan; Department of Hospital Pharmacy & Pharmacology, Asahikawa Medical University, Asahikawa, Hokkaido, Japan

**Keywords:** trabecular meshwork, shear stress, aqueous humor, extracellular matrix, glaucoma, intraocular pressure

## Abstract

**Purpose:** To investigate the changes in extracellular matrix (ECM) gene and protein expressions in human trabecular meshwork (HTM) cells in response to mechanical fluid flow stimulation.

**Methods:** HTM cells were cultured on a glass plate and exposed to shear stress (0, 0.2, and 1.0 dyne/cm^2^) for 12 hours. Changes in gene expressions were evaluated using microarray analysis. The representative genes related to ECM metabolism underwent real-time reverse-transcriptase polymerase chain reaction. Fibronectin (FN) and collagen (COL) IV levels in the supernatant were evaluated using immunoassays. Rho-associated coiled-coil-containing protein kinase (ROCK) activity also was investigated.

**Results:** After stimulation, transforming growth factor β2 mRNA levels were significantly (p < 0.01) lower than that of the static control (0 dyne/cm2 for 12 hours). Matrix metalloproteinase 2 mRNA levels were significantly (p < 0.05) higher than the static control. COL type 1 alpha 2 mRNA, COL type 4 alpha 2 mRNA, and COL type 6 alpha 1 mRNA levels were significantly (p < 0.05, < 0.01, and < 0.05, respectively) higher than the static control. The mean ± standard deviation FN levels (ng/mL) in the supernatant after stimulation (0, 0.2, 1.0 dyne/cm^2^) were 193.7 ± 7.6, 51.5 ± 21.8, and 34.9 ± 23.6, respectively (p < 0.01). The FN and COL IV levels and ROCK activity were significantly (p < 0.01 and < 0.05, respectively) lower than the static control.

**Conclusions:** Changes in gene and protein expressions related to ECM metabolism occurred in HTM cells after stimulation. Specifically, the suppression of FN and COL IV production might explain the importance of mechanical fluid flow stimulation on maintenance of the normal aqueous humor outflow.

## Introduction

Intraocular pressure (IOP) is maintained through a proper function of the aqueous humor, which is produced by the ciliary body [1]. About 70% to 95% of the aqueous humor drains through the conventional outflow pathway [2]. Therefore, normal function of this outflow component is important to the IOP homeostasis and prevention of glaucoma [3]. Increased aqueous outflow resistance in this component is the main cause of glaucoma accompanied by elevated IOP [4, 5].

The trabecular meshwork (TM), juxtacanalicular meshwork, and Schlemm’s canal, the collector channels, and the episcleral veins comprise conventional outflow pathway. Among those, extracellular matrix (ECM) in TM tissue, which are composed of collagen (COL) or fibronectin (FN) [6], is critical for the homeostatic maintenance of the normal outflow resistance [7]. Of note, recent studies revealed that ECM turnover is regulated by mechanical stress, at least in part [8-10].

Experimentally, mechanical stretching to TM cells upregulates gelatinase A and membrane type-1 matrix metalloproteinase (MMP) and reciprocally downregulates tissue inhibitors of metalloproteinases (TIMP)-2 [11]. Additionally, mechanical stretching by pulsatile IOP decreases outflow facility of both human and porcine anterior segments [12]. Moreover, cyclic mechanical stress alters the contractility of the TM cells [13]. Mechanistically, the mechanosensitive receptors on the TM cell surface reportedly sense the deformation of the ECM resulting from the IOP fluctuations via integrin-based cell-matrix contact [14, 15]. Collectively, these findings suggest that mechanical stress applied to the TM cells is important for the ECM turnover.

Clinically, it is generally considered that the TM function (e.g., ECM synthesis, phagocytosis, contractility) deteriorates after glaucoma filtration surgery or development of peripheral anterior synechia over a long period. In addition, the aqueous outflow to the TM after filtration surgery decreases to about 10% of the preoperative level [16], because most of the aqueous humor drains thorough trabeculectomy hole to the filtering bleb, fueling speculation that the mechanical fluid flow stimulation by aqueous outflow to the TM also has an important role in maintaining the TM function. However, to our knowledge, no previous reports have focused on the effect of mechanical fluid flow stimulation on TM cells and its effect on ECM turnover, although the effects of mechanical stimulation by stretching on TM cells have been previously investigated [8-11].

In the current study, to elucidate the role of mechanical fluid flow stimulation on the ECM metabolism in the TM, we applied mechanical fluid flow stimulation to the human trabecular meshwork (HTM) cells and investigated the gene and protein changes related to the ECM metabolism. In addition, because Rho-associated coiled-coil-containing protein kinase (ROCK) is related to the regulation of ECM component expression [17], the ROCK activity also was investigated.

## Materials and methods

### Cell culture

The primary HTM cells obtained from ScienCell (Cat. No. 6590, Lot No. 3423, Carlsbad, CA, USA) were maintained in Fibroblast Medium (ScienCell) containing 5% fetal bovine serum and fibroblast growth supplements and penicillin/streptomycin (FGS, P/S solution, ScienCell), according to the manufacturer’s protocol. The HTM cells were cultured at the bottom of poly-L-lysine-coated flasks (2 μg/cm^2^). The cultured cells were passaged with 0.05% trypsin 2 mM ethylenediaminetetraacetic acid solution and seeded on glass plates (70 × 100 × 1.3 mm) (Matsunami Glass, Kishiwada, Japan) coated with 0.02% type I collagen (Angio-Proteomie, Boston, MA, USA). We used the HTM cells from between passages 5 and 7. The current study was performed in accordance with the tenets of the Declaration of Helsinki.

### Shear stress experiments

Shear stress was applied to the confluent HTM cells using a parallel plate-type flow chamber. We previously described the shear stress experiments [18-20]. Briefly, this flow circuit included a flow chamber, a peristaltic pump (SJ1220, ATTO, Tokyo, Japan), and a medium reservoir connected using silicone tubes. The culture medium was circulated continually with a peristaltic pump in an incubator at 37°C with 5% carbon dioxide. The shear stress inside the flow chamber was calculated by the formula, τ = μ·6Q/a^2^b, where τ is the shear stress (dyne/cm^2^), μ is the viscosity of the perfused fluid (0.00796 poise), Q the volumetric flow rate (mL/s), a the flow channel height (0.04 cm), and b the flow channel width (5.5 cm) in the cross section. To generate shear stress using a perfused medium, we used Dulbecco’s Modified Eagle’s medium (Wako Pure Chemical Industries, Ltd., Osaka, Japan) supplemented with 100 U/mL penicillin and 100 μg/mL streptomycin sulfate and GlutaMAX-I supplement (Life Technologies, Carlsbad, CA, USA) without serum. The medium viscosities were measured using a viscometer (ViscoLab 4000, Japan Controls, Tokyo, Japan). In the current study, the HTM cells were exposed to the following magnitudes of laminar shear stress: 0 dyne/cm^2^ (static control) and 0.2 and 1.0 dyne/cm^2^. After exposure to fluid flow for 12 hours, the glass plates on which the HTM cells were cultured were removed from the flow chamber and rinsed in phosphate buffered saline (PBS), and the HTM cells were collected

### Microarray analysis for gene expressions

A DNA microarray assay was performed by Hokkaido System Science (Sapporo, Japan). Briefly, the total RNA from the HTM cells was amplified and transcribed into fluorescent cRNA using a Low Input Quick Amp Labeling Kit, One Color (Agilent Technologies, Santa Clara, CA, USA). The labeled cRNA was purified using an RNeasy Mini Spin Kit (QIAGEN, Germantown, MD, USA) and hybridized to the human microarray kit (SurePrint G3 Human GE 8 × 60 K version 3.0, Agilent Technologies). The raw data were normalized using a Gene Spring Software (version 13.0, Agilent Technologies). The normalized microarray data then were compared with the real-time RT-PCR analysis.

### Real-time RT-PCR analysis for representative genes related to ECM metabolism

The HTM cells were collected with a scraper, and the total RNA was extracted using a NucleoSpin RNA kit (Takara, Shiga, Japan). The total RNA (25 μg/mL) underwent reverse transcription using a Transcriptor First Strand cDNA Synthesis Kit (Roche, Basel, Switzerland), according to the manufacturer’s instructions, after which real-time PCR was performed using the Universal ProbeLibrary and Light Cycler 480 (Roche). The specific primer pairs are shown in Table 1. For all amplifications, the cycling conditions were as follows: an initial denaturation period for 5 minutes at 95°C, followed by 45 cycles of 10 seconds at 95°C, 30 seconds at 60°C, and 1 second at 72°C. The quantification of each gene expression signal was normalized with respect to the signal for the *glucose-6-phosphate dehydrogenase* (*G6PDH*) gene. The relative fold changes in the expression of each gene were determined using the 2^-ΔΔCt^ method.

**Table 1.**
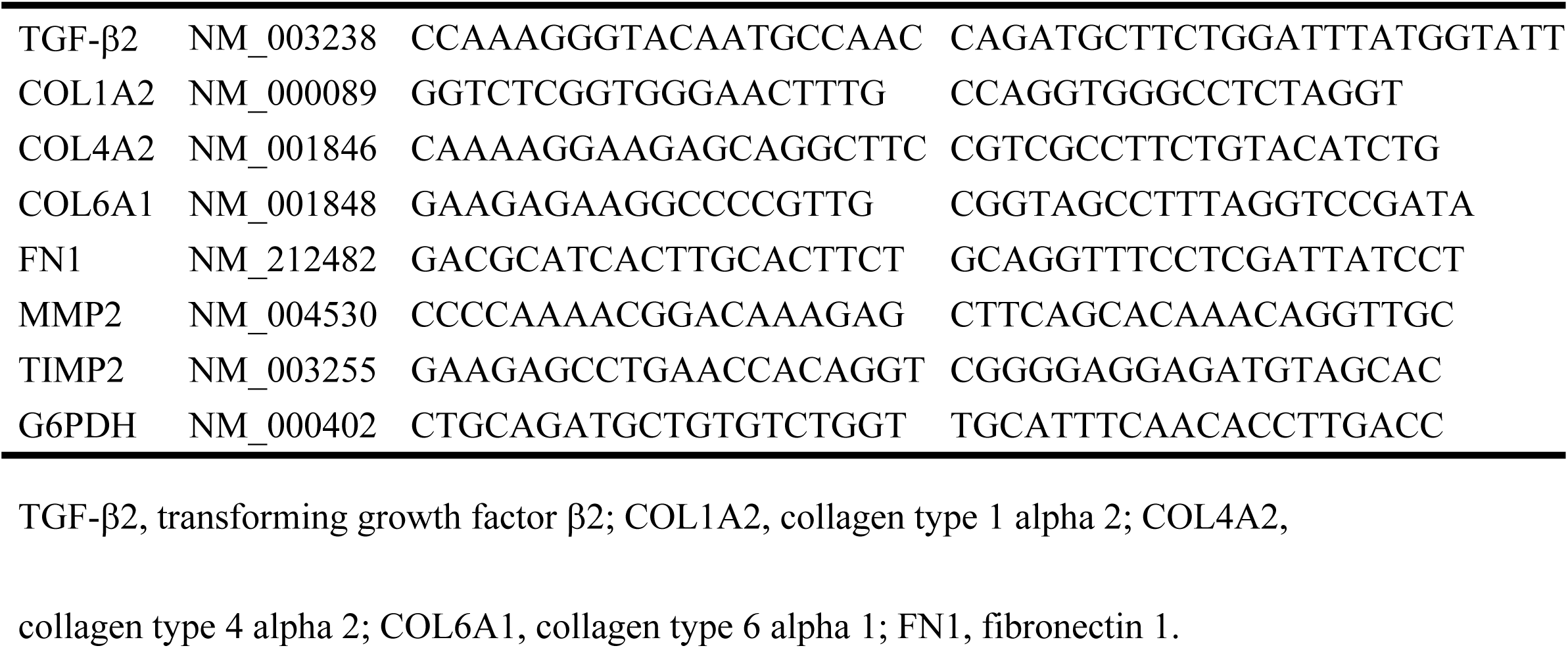
Primer Sequences Used for Gene Expression Analysis by Real-Time RT-PCR.

### Enzyme-linked immunosorbent assay (ELISA) for detecting FN levels

FN in the culture medium supernatant was measured using an ELISA kit (R&D Systems, Minneapolis, MN, USA), according to the manufacturer’s instructions. Culture medium supernatants were collected from the HTM cells with or without exposure to shear stress for 12 hours. The optical density of the samples and fibronectin standard were determined using a microplate reader.

### Western blot analysis for detection of FN and COL IV levels

The HTM cells were washed with PBS and lysed with RIPA Lysis Buffer (Merck Millipore, Darmstadt, Germany) containing protease inhibitor cocktail tablets (Roche) and phosphatase inhibitor cocktail tablets (Roche) and phenylmethylsulfonyl fluoride. The lysates then were centrifuged at 13,000 rpm for 20 min at 4°C, and the resultant supernatants were collected. The total protein concentration was measured using a NanoDrop Fluorospectrometer (Thermo Fisher Scientific, Waltham, MA, USA). Equal amounts of protein were loaded into each well and separated by sodium dodecyl sulfate polyacrylamide gel electrophoresis and subsequently transferred to nitrocellulose membranes by electroblotting. The membranes were blocked with PVDF Blocking Reagent (Toyobo, Osaka, Japan) and incubated in Can Get Signal (Toyobo) containing the following antibodies for 1 hour: monoclonal β-actin antibodies (mouse, #3700, 1:2000 dilution) (Cell Signaling Technology, Danvers, MA, USA), polyclonal FN antibodies (rabbit, F3648, 1:1000 dilution), (Sigma-Aldrich, St. Louis, MO, USA), polyclonal COL IV antibodies (rabbit, SAB4500369, 1:1000 dilution) (Sigma-Aldrich), and horseradish peroxidase-conjugated anti-rabbit or anti-mouse IgG secondary antibody (#7076 and #7074, 1:10000 dilution) (Cell Signaling Technology). The membranes were exposed to ECL Prime Western Blotting Detection Reagent (GE Healthcare, Piscataway, NJ, USA) and examined using LAS-3000 Imager (Fujifilm, Tokyo, Japan).

### ROCK activity analysis

The cell lysates were collected from the HTM cells with or without exposure to shear stress for 12 hours. The total protein concentration was measured using a NanoDrop Fluorospectrometer. Equal amounts of protein then were measured by an ELISA using a 96-well ROCK Activity Assay Kit (Cell Biolabs, San Diego, CA), according to the manufacturer’s instructions. Absorbance were measured on a microplate reader at 450 nm and results were compared to the static control.

### Statistical analysis

The data were analyzed using GraphPad Prism version 5 (GraphPad Software, San Diego, CA). Quantitative data were analyzed using the Dunn’s nonparametric comparison for post-hoc testing after the Kruskal-Wallis test. P < 0.05 was considered significant.

## Results

### Gene expressions in HTM cells in response to shear tress

Our microarray analysis investigated 62,976 genes. Among those, we focused on representative genes related to ECM metabolism and compared the gene expression levels of the static control to those exposed to shear stress (0.2 dyne/cm^2^) for 12 hours. The gene categories, names, and fold changes are shown in Table 2. Those genes listed then were analyzed by real-time RT-PCR to ascertain the results obtained by microarray analysis. As a result, transforming growth factor (TGF)-β2 mRNA levels were significantly lower than that of the static control (0.2 dyne/cm^2^, 0.65-fold vs. static control, p < 0.01) (Fig. 1). The MMP2 mRNA levels were significantly higher (0.2 dyne/cm^2^, 2.35-fold vs. static control, p < 0.05 and 1.0 dyne/cm^2^, 2.96-fold vs. static control, p < 0.05) (Fig. 2A), while the differences in the TIMP2 mRNA levels were not significant (0.2 dyne/cm^2^, 1.49-fold vs. static control, and 1.0 dyne/cm^2^, 2.03-fold vs. static control, p = 0.055) (Fig. 2B) The COL type 1 alpha 2 (COL1A2) mRNA (1.0 dyne/cm^2^, 1.79-fold vs. static control, p < 0.05), COL type 4 alpha 2 (COL4A2) mRNA (1.0 dyne/cm^2^, 3.11-fold vs. static control, p< 0.01), and COL type 6 alpha 1 (COL6A1) mRNA (1.0 dyne/cm^2^, 1.91-fold vs. static control, p<0.05) levels were significantly higher than the static control. Although the FN1 mRNA levels were higher than the static control (1.0 dyne/cm^2^, 1.87-fold vs. static control), the differences were not significant (p = 0.085) (Fig. 3).

**Table 2.** Comparison of Gene Expression Levels of the Static Control with Those Exposed to Shear Stress (0.2 dyne/cm^2^) for 12 Hours.

| Gene Category | Gene Name (symbol) | Accession Number | Fold Change (vs. static control) |
| --- | --- | --- | --- |
| TGF-β family | TGF-β1 | NM_000660 | 1.71 |
| | TGF-β2 | NM_003238 | 0.52 |
| | TGF beta receptor 2 (TGFβR2) | NM_001024847 | 0.60 |
|  | Bone morphogenetic protein 5 (BMP5) | NM_021073 | 0.29 |
|  | BMP6 | NM_001718 | 3.49 |
| ECM | COL1A2 | NM_000089 | 0.92 |
|  | COL2A1 | NM_001844 | 1.05 |
|  | COL3A1 | NM_000090 | 0.62 |
|  | COL4A2 | NM_001846 | 1.47 |
|  | COL5A2 | NM_000093 | 0.74 |
|  | COL6A1 | NM_001848 | 1.00 |
|  | FN1 | NM_054034 | 1.41 |
|  | Elastin | NM_001278939 | 2.07 |
| | Laminin β3 | NM_001017402 | 2.27 |
| ECM remodeling | MMP1 | NM_002421 | 3.01 |
|  | MMP2 | NM_004530 | 2.24 |
|  | MMP3 | NM_002422 | 3.26 |
|  | MMP9 | NM_004994 | 1.23 |
|  | TIMP1 | NM_003254 | 1.04 |
|  | TIMP2 | NM_003255 | 0.99 |
|  | Plasminogen activator inhibitor type 1 | NM_000602 | 11.91 |
|  | Connective tissue growth factor | NM_001901 | 1.99 |

**Fig. 1.**
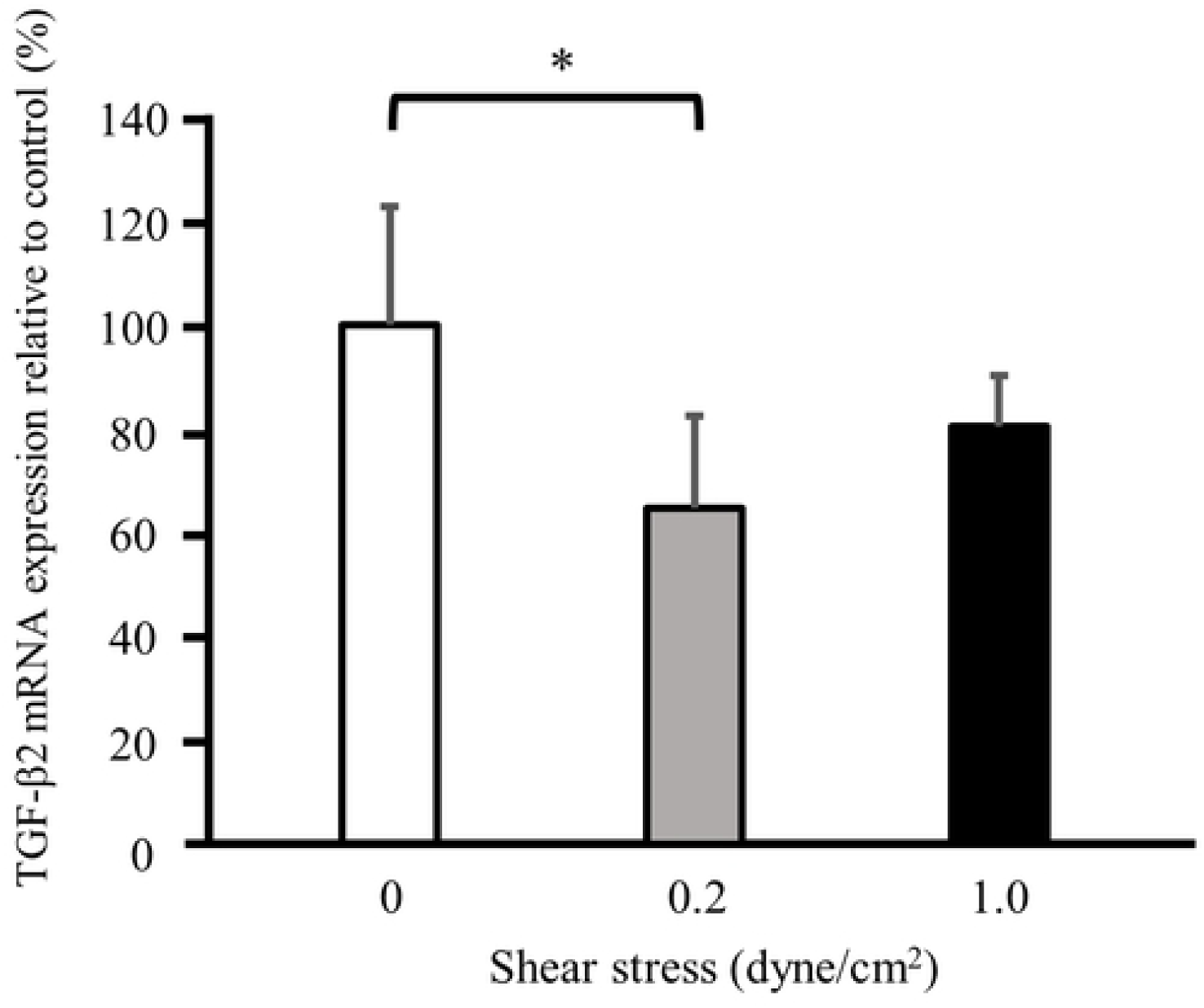
Real-time RT-PCR analysis of TGF-β2 mRNA expression in HTM cells. TGF-β2 mRNA levels in the HTM cells exposed to shear stress (0.2 dyne/cm^2^) for 12 hours are significantly lower than the static control. The data are expressed as the means ± standard deviation (n = 5). *p < 0.01.

**Fig. 2.**
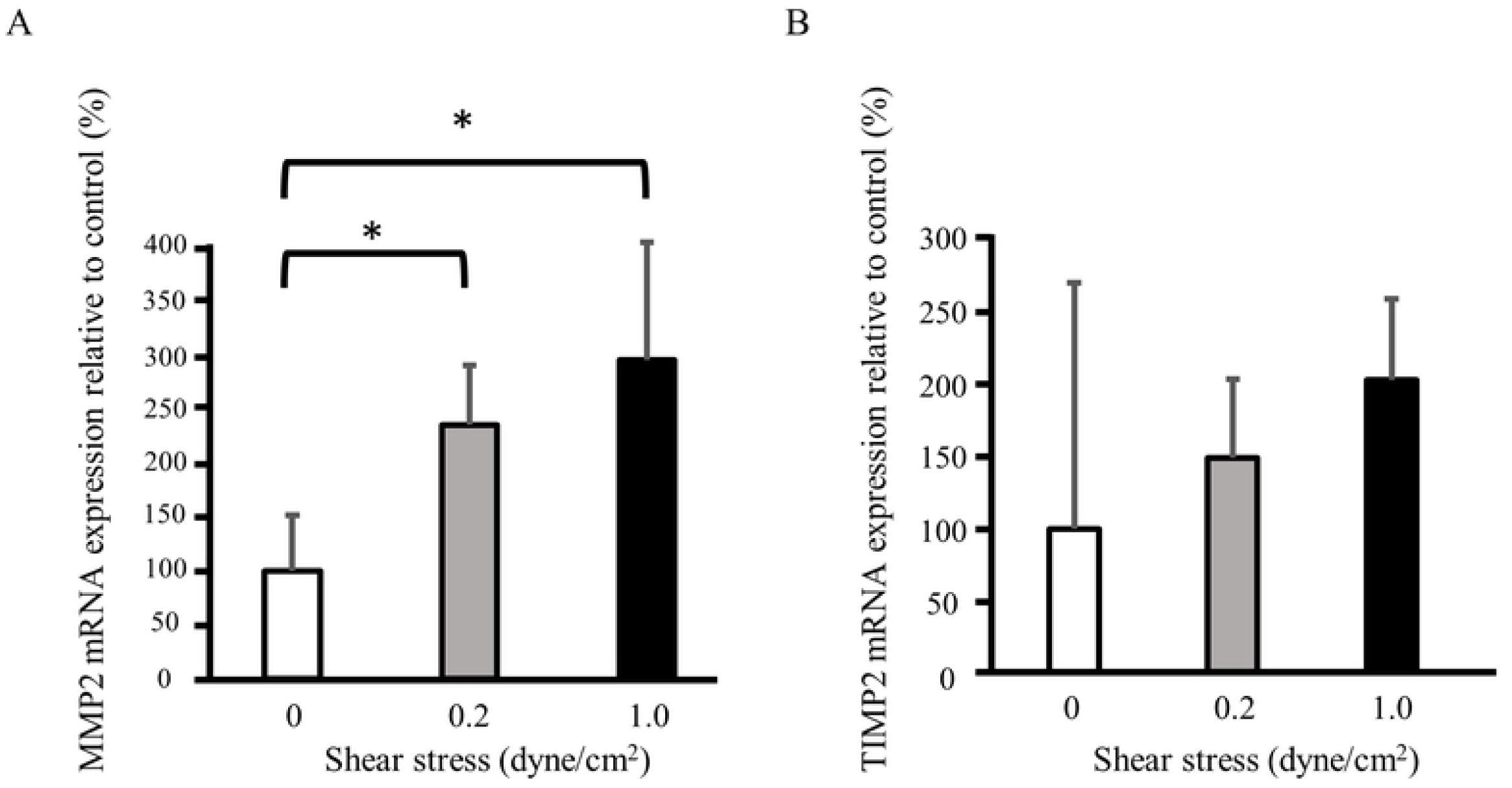
Real-time RT-PCR analysis of MMP2 mRNA and TIMP2 mRNA expression in HTM cells. (A) MMP2 mRNA levels in the HTM cells exposed to shear stress (0.2 and 1 dyne/cm^2^) for 12 hours are significantly higher than the static control. (B) Although TIMP2 mRNA levels are higher than the static control after stimulation, the differences are not significant. The data are expressed as the means ± standard deviation (n = 5). *p < 0.05.

**Fig. 3.**
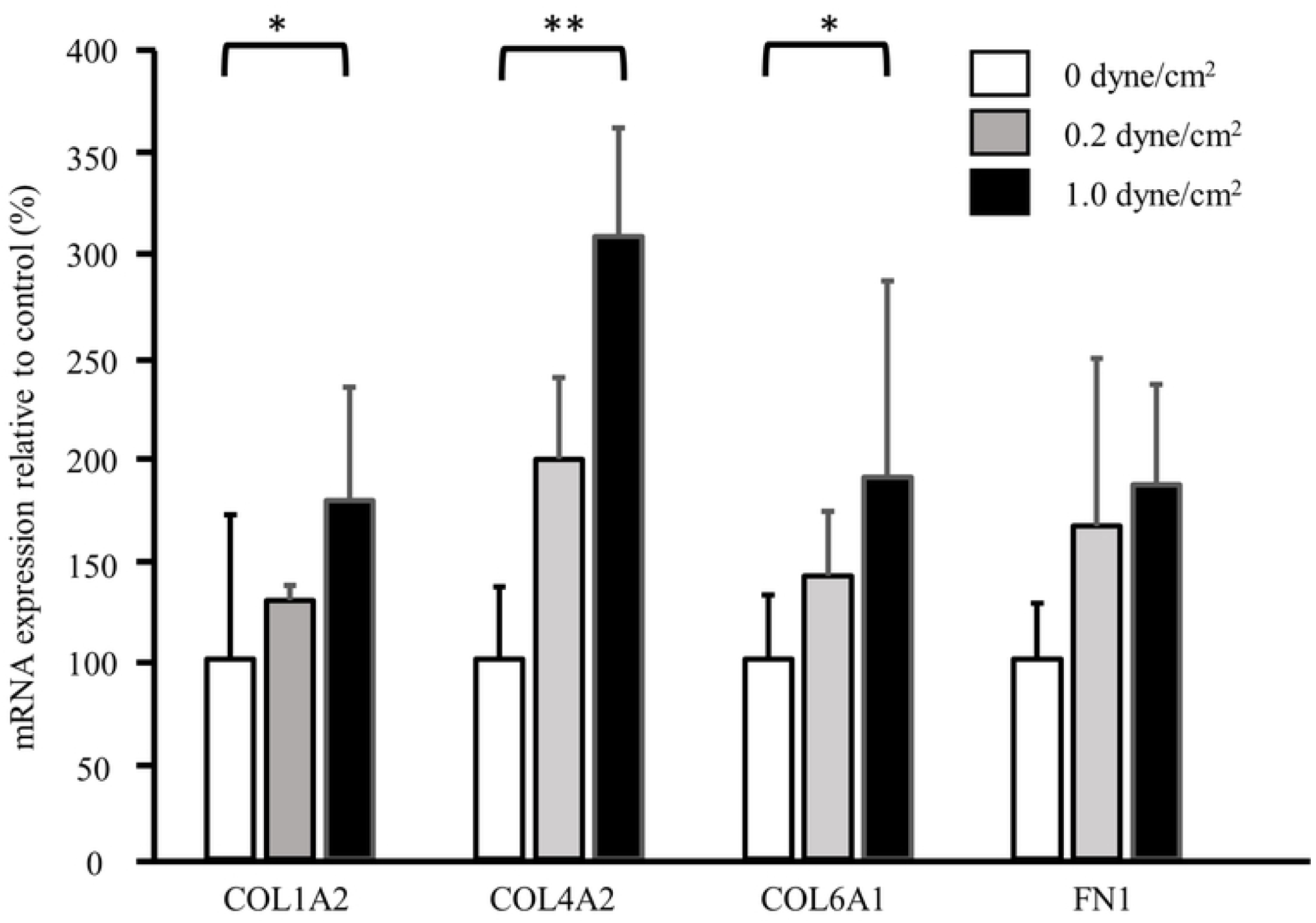
Real-time RT-PCR analysis of COL1A2, COL4A2, COL6A1, and FN1 mRNA expression in HTM cells. COL1A2, COL4A2, and COL6A1 mRNA levels in the HTM cells exposed to shear stress (1 dyne /cm^2^) for 12 hours are significantly higher than the static control. Although the FN1 mRNA levels are higher than the static control after stimulation, the differences are not significant. The data are expressed as the means ± standard deviation (n = 4). *p < 0.05 and **p < 0.01.

### FN and COL IV levels in the supernatant after exposure to shear stress

The mean (± standard deviation) FN levels (ng/mL) in the medium exposed to shear stress (dyne/cm^2^) of 0, 0.2, and 1.0 for 1 hour were 31.6 ± 19.6, 30.0 ± 18.7, and 29.5 ± 13.6, respectively. The values for 12 hours were 193.7 ± 7.6, 51.5 ± 21.8, and 34.9 ± 23.6, respectively. After 12 hours, the FN levels were significantly (p < 0.01) lower at 1.0 dyne/cm^2^ than the static control (Fig. 4). Representative images of each band obtained by Western blot analysis are shown in Fig. 5A. The FN levels were significantly lower at 0.2 dyne/cm^2^ (p < 0.05) and 1.0 dyne/cm^2^ (p < 0.01) than the static control. The COL IV levels at 1.0 dyne/cm^2^ also were significantly (p < 0.01) lower than the static control (Fig. 5B).

**Fig. 4.**
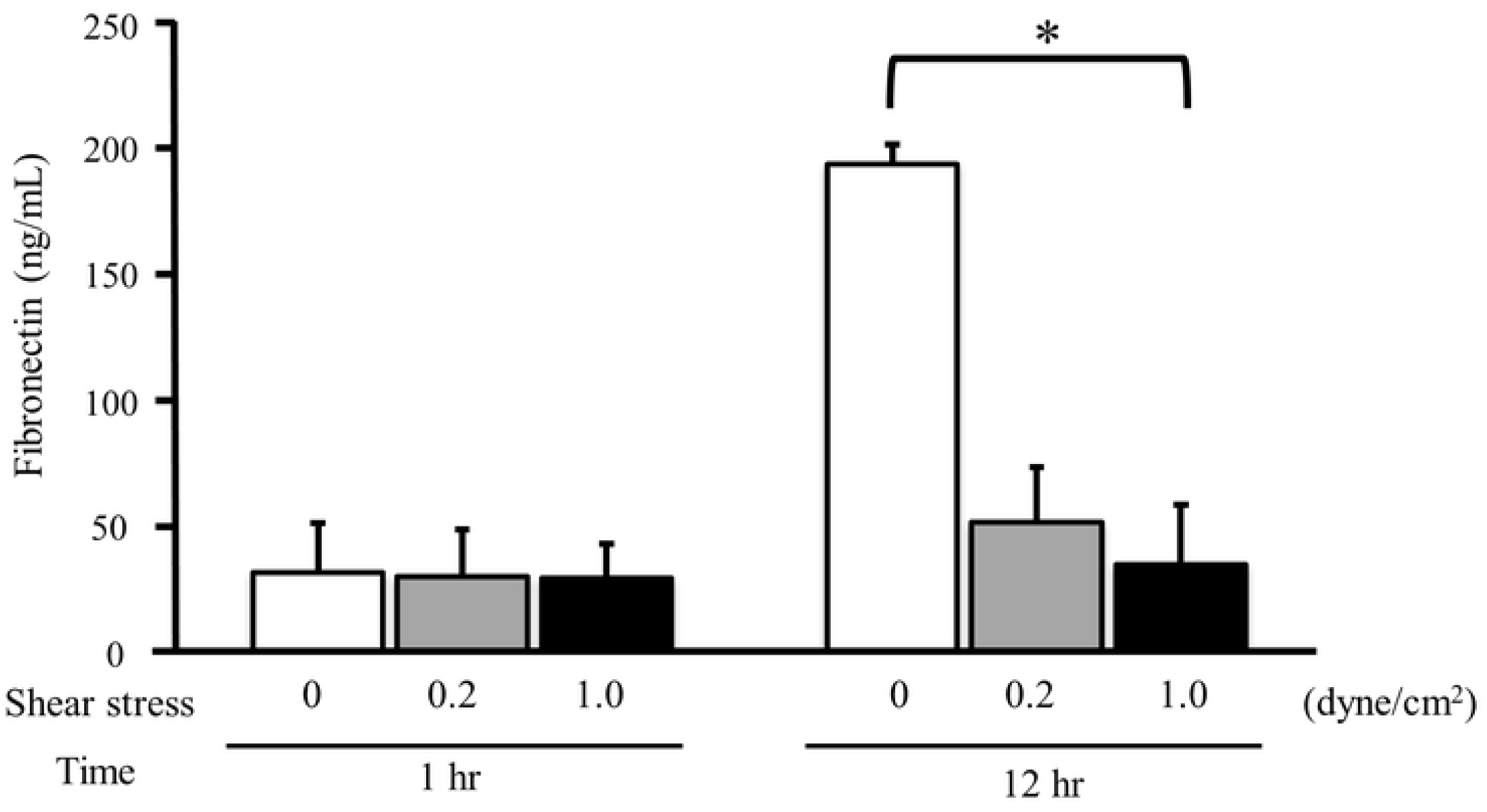
The concentration of FN in the culture supernatant after exposure to shear stress. After exposure to shear stress (1 dyne /cm^2^) for 12 hours, the levels of FN are significantly lower than the static control. The data are expressed as the means ± standard deviation (n = 5). **p < 0.01. hr, hours.

**Fig. 5.**
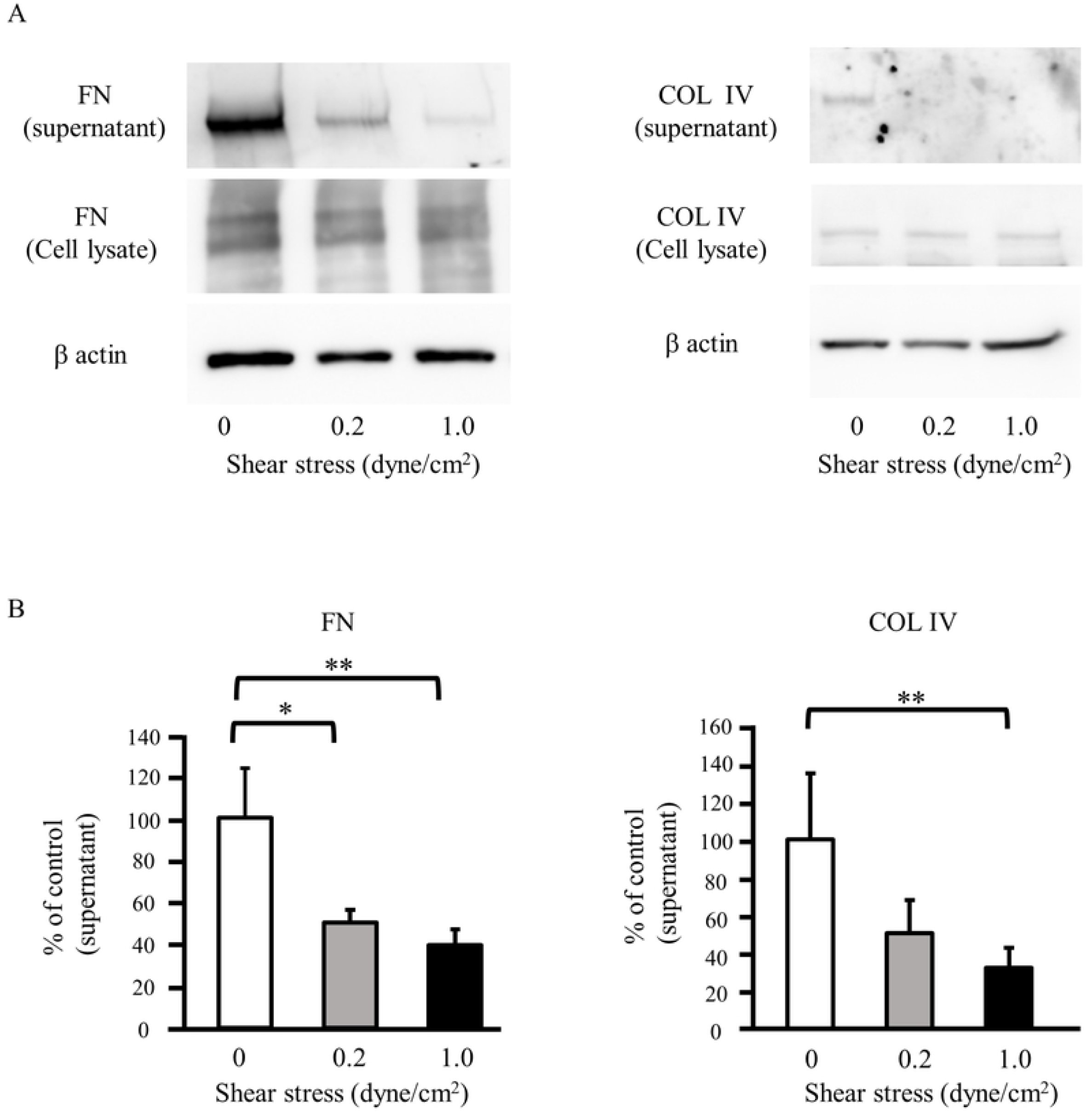
FN and COL IV levels in HTM cells after exposure to shear stress. (A) Representative images of Western blot analysis of FN and COL IV expression. (B) Quantitative assessment of the intensity of each band determined by densitometry. The FN and COL IV levels are significantly lower compared to the static control after stimulation. The data are expressed as the means ± standard deviation (FN, n = 6; COL IV, n = 5). *p < 0.05 and **p < 0.01.

### ROCK activity in HTM cells after exposure to shear stress

To examine the ROCK activity, we carried out the ELISA technique. Our experiment showed that the ROCK activity was significantly lower than the static control after exposure to shear stress (0.2 dyne/cm^2^, 0.78-fold vs. static control, p < 0.05) (Fig. 6).

**Fig. 6.**
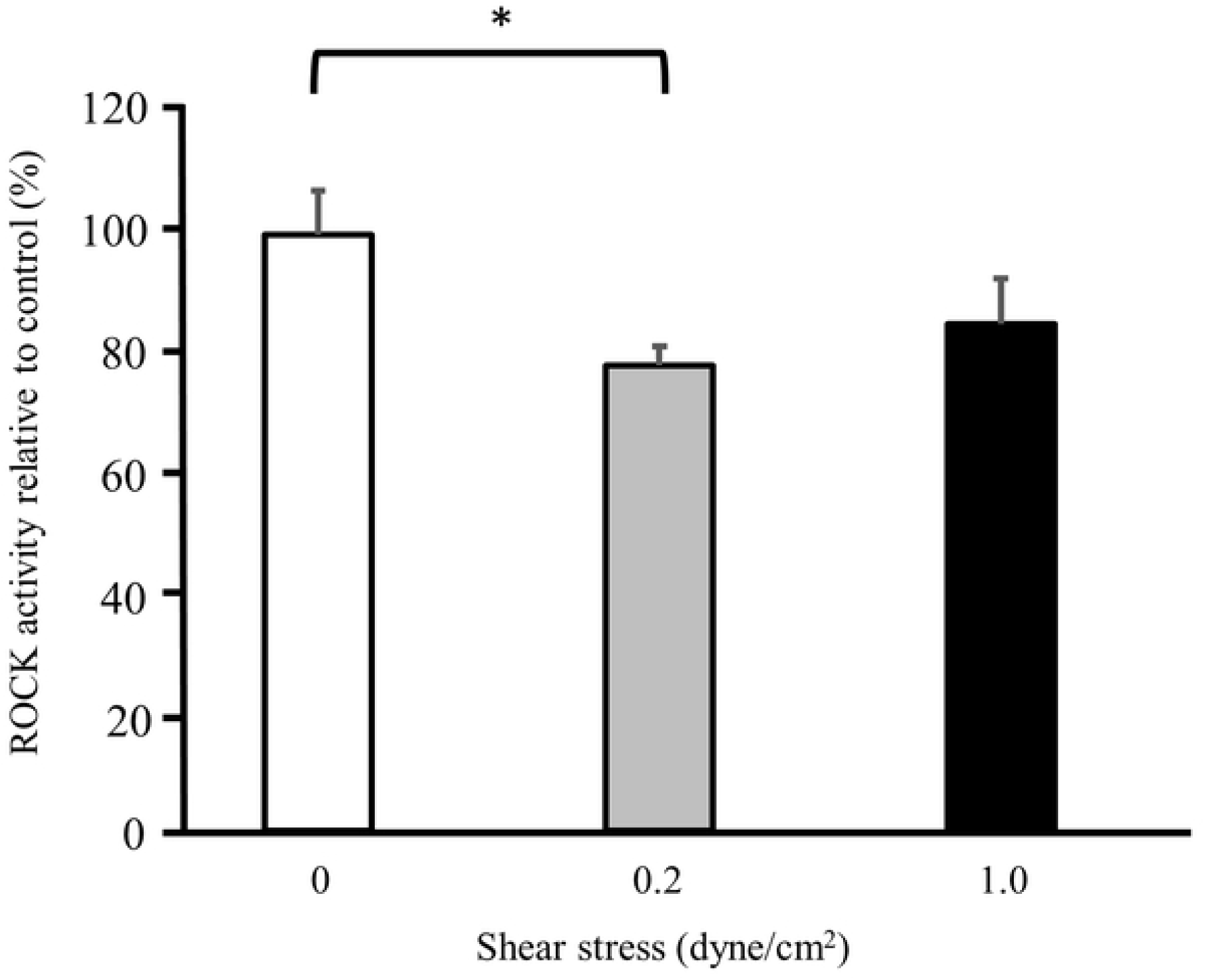
ROCK activity in HTM cells after exposure to shear stress. The ROCK activity is significantly lower than the static control after exposure to shear stress (0.2 dynes/cm^2^) for 12 hours. The data are expressed as means ± standard deviation (n = 4). *p < 0.05.

## Discussion

In the current study, we applied shear stress to HTM cells and investigated the changes in gene or protein expressions related to ECM metabolism. Because FN, laminin, and COL IV are the main ECM components of the JCT [6], we focused on those components in our experiments. As a result, FN and COL IV levels in the supernatant, in which HTM cells were exposed to shear stress for sufficient time, were significantly lower than the static control. This implies that FN and COL IV production by HTM cells are suppressed in the presence of shear stress. Our microarray analysis and subsequent real-time RT-PCR also showed lower levels of TGF-β2 mRNA and elevated levels of MMP2 mRNA expression with increased shear stress. Because down-regulation of TGF-β2 and the increased MMP2 promote ECM degradation, HTM cells may promote degradation of the ECM in the presence of shear stress. Regarding the increases in COL mRNA and FN1 mRNA, we considered these as compensatory responses (i.e., negative feedback mechanism). Our results agreed with previous reports that examined the effects of mechanical stimulation by stretching [21, 22] or IOP pulsation on TM cells in that the mechanical stimulation might have an important role in ECM turnover in the TM [12].

Previously, fluid flow stress was reported to affect the pathological condition or homeostasis in some ocular tissues, e.g., flow stress on cultured human corneal epithelial cells affected TGF-β signaling, which may be associated with a delayed wound-healing mechanism during blinking [20]. Flow stress also can cause corneal endothelial cell damage or loss after laser iridotomy [23] or gene expression of endovascular cells, which may contribute to the vasoregulatory and antithrombotic properties of the retinal vessels [18]. Although the current study focused on mechanical stimulation of the fluid flow on the HTM cells, a previous report also investigated the responses of HTM cells exposed to shear stress. Ashpole et al. [24] reported that when shear stress (10 dyne/cm^2^) was applied to human SC endothelial (SCE) cells, they responded similarly to vascular endothelial cells, i.e., shear stress also triggered nitric oxide (NO) production on human SCE cells. Those authors concluded that increased shear stress to SCE cells during SC collapse in the presence of elevated IOP may partly mediate IOP homeostasis by NO production. Although, in that study, they also investigated the responses of HTM cells exposed to a shear stress (10 dyne/cm^2^), they only focused on NO production. In the current study, we applied shear stress to HTM cells, but the strength was very weak (0.2 or 1.0 dyne/cm^2^). Therefore, the current study differed from their research in that we focused on the gene or protein changes related to ECM metabolism in response to very weak shear stress.

In our microarray analysis, we focused on the gene changes related to the TGF-β superfamily and ECM components and remodeling. As a result, the expression of TGF-β1 mRNA was observed, although it could not be confirmed by subsequent real-time RT-PCR in our experiments. A previous study reported increased production of TGF-β1 after cyclic mechanical stress in the TM [9]. Further, the increased expression of connective tissue growth factor (CTGF) mRNA and plasminogen activator inhibitor-1 mRNA also were observed, suggesting changes in many genes involved in ECM metabolism when share stress was applied to HTM cells. The simultaneous increase of TGF-β1 mRNA and CTGF mRNA may be reasonable, because CTGF is also up-regulated by TGF-β1 [25]. Regarding bone morphogenetic protein (BMP) 5, which was down-regulated in the current study, it is expressed in the TM and is involved in the normal formation and function of the TM [26]. Although the role of BMP6 in the TM has not been yet elucidated, altered expression of members of the BMP family may cause functional changes in the TM [27].

In the current study, the ROCK activity in the HTM cells after stimulation was lower than the static control. Because the fibrogenic effect caused by TGF-β2 is mediated by activating ROCK, the decreased ROCK activity may contribute partly to decreased production of FN and COL IV. The recently launched anti-glaucoma drug ROCK inhibitor (ripasudil) acts on the conventional outflow pathway and decreases IOP by increasing aqueous humor outflow through diverse mechanisms [28, 29]. Among the actions of ripasudil on the TM-SC pathway, the inhibition of ECM production [30] and the suppression of TM cell contraction were considered as the functions targeting the JCT [31]. Given the current results, improvement of the aqueous humor outflow through the TM by ripasudil may further promote the ECM metabolism, contributing to further decreases in IOP when the drug is used over the long term.

Although, in the current study, the HTM cells were stimulated by steady fluid flow using our shear stress experimental system, it is presumed that the fluid flow in the TM is probably not a laminar or steady flow in normal eyes. A previous report that simulated the wall shear stress in each component of the conventional outflow pathway reported that the shear stress on the inner wall of the SC was about 0.01 dyne/cm^2^ [32]. Therefore, in the JCT, which is the principal tissue of the ECM metabolism in the TM, the shear stress might be equal to or less than that in the inner wall of the SC (i.e., much weaker stimulation than the current experiment). Further, it also has been reported that the TM tissue pulsates in conjunction with the heart rate [33, 34]. Taken together, considering this histologic feature of the meshwork, the aqueous outflow in the TM may be turbulent. Further investigation is needed to determine if the results of the current shear stress experiment also occur in vivo.

In conclusion, gene and protein changes related to ECM metabolism were observed as a result of culturing of the HTM cells in the presence of shear stress. The current results suggested that the mechanical stimulation of aqueous fluid flow to the TM promotes ECM turnover, contributing to maintenance of IOP homeostasis in normal eyes.

## Acknowledgments

We are grateful to C. Matsumoto and A. Tanner for their excellent technical assistance.

## Author contributions

Conceived and designed the experiments: KY, MK, TU, YT, AI. Performed the experiments: KY. Analyzed the data: KY, MK, YS, MU. Contributed reagents/materials/analysis tools: MK, TU, AI, AY. Wrote the paper: KY, MK.

